# Humans can use positive and negative spectrotemporal correlations to detect rising and falling pitch

**DOI:** 10.1101/2024.08.03.606481

**Authors:** Parisa A. Vaziri, Samuel D. McDougle, Damon A. Clark

## Abstract

To discern speech or appreciate music, the human auditory system detects how pitch increases or decreases over time. However, the algorithms used to detect changes in pitch, or pitch motion, are incompletely understood. Here, using psychophysics, computational modeling, functional neuroimaging, and analysis of recorded speech, we ask if humans can detect pitch motion using computations analogous to those used by the visual system. We adapted stimuli from studies of vision to create novel auditory correlated noise stimuli that elicited robust pitch motion percepts. Crucially, these stimuli are inharmonic and possess no persistent features across frequency or time, but do possess positive or negative local spectrotemporal correlations in intensity. In psychophysical experiments, we found clear evidence that humans can judge pitch direction based only on positive or negative spectrotemporal intensity correlations. The key behavioral result—robust sensitivity to the negative spectrotemporal correlations—is a direct analogue of illusory “reverse-phi” motion in vision, and thus constitutes a new auditory illusion. Our behavioral results and computational modeling led us to hypothesize that human auditory processing may employ pitch direction opponency. fMRI measurements in auditory cortex supported this hypothesis. To link our psychophysical findings to real-world pitch perception, we analyzed recordings of English and Mandarin speech and found that pitch direction was robustly signaled by both positive and negative spectrotemporal correlations, suggesting that sensitivity to both types of correlations confers ecological benefits. Overall, this work reveals how motion detection algorithms sensitive to local correlations are deployed by the central nervous system across disparate modalities (vision and audition) and dimensions (space and frequency).

## Introduction

From discriminating phonemes to being moved by Bach’s *Partitas*, detecting relative changes in pitch over time is fundamental to human audition, allowing us to perceptually characterize sounds proceeding from low frequency to high frequency and vice versa (**Fig. 1a**). Indeed, in everyday speech we use both intonation and lexical tones — including complex rising and falling pitches — to signify meaning (Hirst and Di Cristo 1998; Gandour 1983; Yip 2002). In English, for instance, rising pitch at the end of a sentence signifies a question. In Mandarin Chinese, changes of pitch within words conveys fundamental differences in meaning. But how does the human auditory system detect changes in pitch?

**Figure 1.**
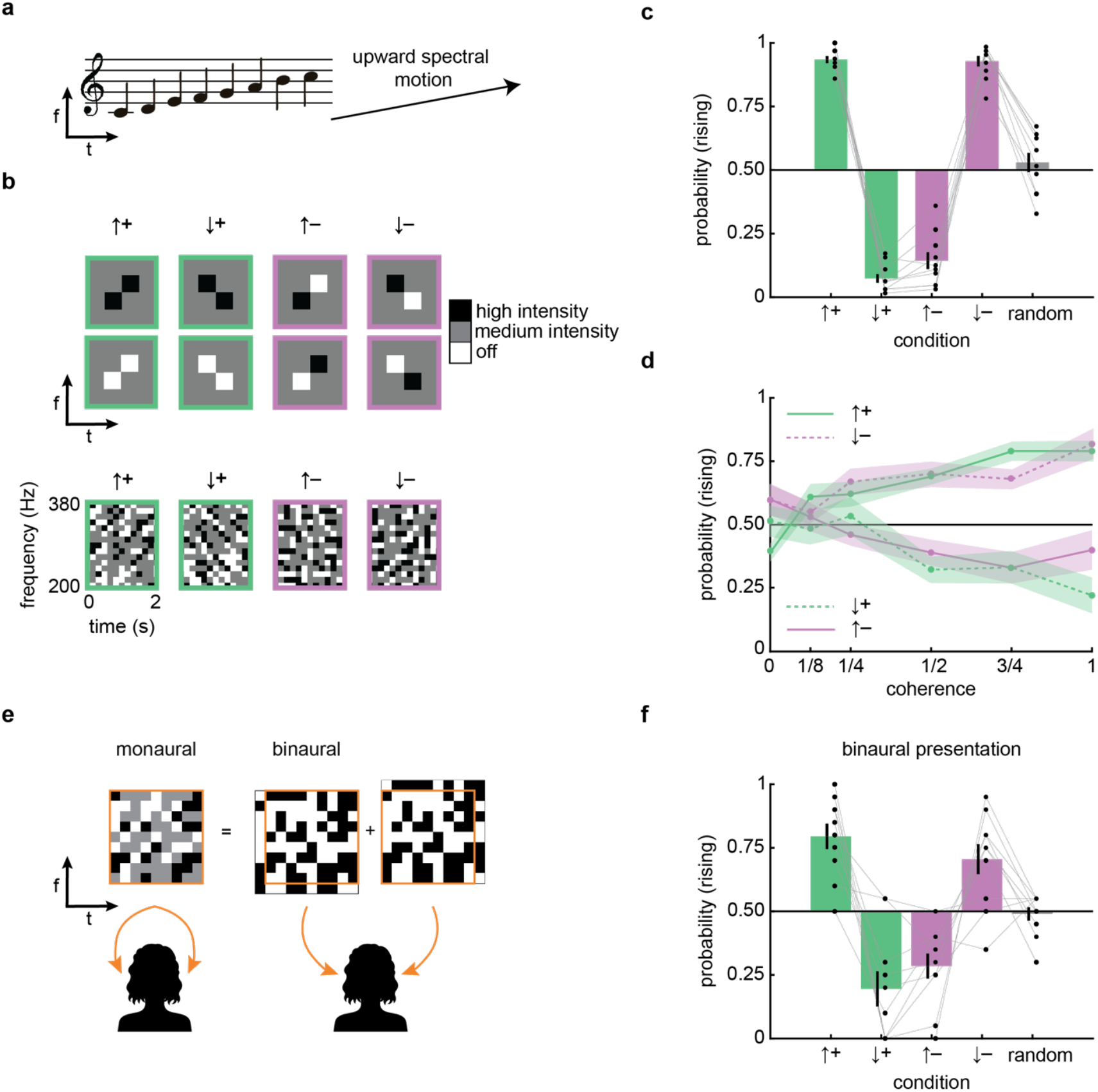
Humans detect auditory motion in pairwise frequency-time correlations. a) Simple schematic of a rising sound written on a music staff and in frequency-time. b) Diagrams showing sample (*top*) and actual (*bottom*) stimuli. Frequency-time correlations can be directed either upward or downward and be either positively or negatively correlated. c) Perceived direction of stimuli with varying direction and correlation. Mean ± SEM over N = 10 subjects. One-sample t-tests revealed significant deviations from chance (0.50) in pitch direction judgements in all four stimulus conditions (all *p*s < 10^−5^). Pitch direction judgements in the random stimulus condition were not significantly different from chance (p = 0.45). Error bars represent mean ± SEM (N = 10). d) Perceived direction of stimuli with varying degrees of correlation (coherence) in the stimulus. The upward-directed positive and downward-directed negative curves were not significantly different (p > 0.05 by a two-way, repeated measures ANOVA); similarly, the downward-directed positive and upward-directed negative curves were also not significantly different (p > 0.05, same test). Both ANOVAs revealed significant main effects of coherence on pitch direction judgements (all *p*s < 10^−5^). Error shading represents ± SEM (N = 10). e) Diagram showing how binaural stimuli were presented to each ear. f) Perceived direction of stimuli with varying directions and correlations using binaural presentation. One-sample t-tests revealed significant deviations from chance (0.50) in pitch direction judgements in all four stimulus conditions (all *p*s < 10^−3^). Pitch direction judgements in the random stimulus condition were not significantly different from chance (p = 0.72). Error bars represent mean ± SEM (N = 10).

Changes in relative pitch can, in principle, be detected in at least two broad ways. The most well-known computation centers on detecting a fundamental frequency (F0), which refers to the lowest frequency in the ladder of harmonics produced by most vibrating objects, including vocal cords. Once F0 is estimated (which is itself not a trivial task (de Cheveigné 1993)), it may be tracked over time to detect change in relative pitch (McDermott and Oxenham 2008). However, relative pitch can also be perceived without detecting a fundamental frequency — psychophysical tests have demonstrated that changes in pitch can be perceived by detecting shifts in multiple constituent frequencies of a sound without needing to compute F0 (McPherson and McDermott 2018). These results indicate that a variety of cues in a sound, not just F0, can support the detection of pitch changes. Importantly, work on such spectral pattern cues typically employs sounds with persistent groupings of tones (Sinnott and Aslin 1985; Demany and Ramos 2005; Aslin 1989; Siedenburg et al. 2023; Demany and Semal 2018) or entire spectra that shift in frequency over time (van der Willigen et al. 2024). These sorts of persistent spectral intensity patterns could in theory be identified and tracked by auditory systems, akin to the detection of unique sound sources in noise (e.g., the cocktail party effect (Cherry 1953; McDermott 2009)), or, more loosely, akin to the detection and tracking of objects in vision. Here, inspired by work in the visual system, we focus on a novel auditory stimulus that possesses no F0 information and, critically, no trackable, persistent features over time (e.g., tonal, timbre-related, etc.), but does contain local positive and negative spectrotemporal correlations in intensity.

Why examine spectrotemporal correlations? In vision, sensing local spatiotemporal correlations is the basis of canonical models for spatial motion detection (Hassenstein and Reichardt 1956; Adelson and Bergen 1985). These models generate sensitivity to pairwise intensity correlations over time and space through a process of linear filtering and nonlinear interactions. Visual sensitivity to intensity correlations can be dramatically revealed by visual illusions, specifically those involving negative spatiotemporal correlations, exemplified by “reverse phi” phenomena (Hassenstein and Reichardt 1956; Anstis and Rogers 1975; Adelson and Bergen 1985). Thus, at least in vision, local intensity correlations serve as an important cue for detecting motion in the environment. These cues feed into local motion detectors that complement parallel systems that detect motion by tracking visual objects over time (Lu et al. 1999; Lu and Sperling 2001; Lu and Sperling 1995). Crucially, the sensitivity to negative correlations in reverse-phi phenomena is fundamentally inconsistent with object or pattern tracking-based models of motion detection.

Existing evidence supports the possibility that local spectrotemporal correlations could aid in the perception of directional pitch changes in audition, without the need for computing F0 or even tracking global spectral patterns. For example, studies of cortical auditory neurons have characterized spectrotemporal receptive fields with shapes that can be oriented over frequency and time (Theunissen et al. 2000; Miller et al. 2002; Depireux et al. 2001). Such receptive fields could support auditory feature tracking, but in principle could also aid in sensing spectrotemporal correlations (Adelson and Bergen 1985). Moreover, one prior study presented a persistent spectral intensity pattern that was repeatedly inverted and frequency-shifted in time, leading to perceptual reversals of pitch direction (Allik et al. 1989). However, the stimulus choice in that study could not isolate negative spectrotemporal correlations, which allow definitive evaluations of the role of correlations in detecting pitch direction. Here, to evaluate how spectotemporal correlations contribute to pitch motion detection, we develop novel auditory stimuli that that lack both persistent spectral features and a common F0, but contain specified local positive and— critically—negative correlations.

## Results

### Spectral motion without features

We set out to test whether humans can detect auditory motion based on local spectrotemporal correlations. To do this, we adapted a stimulus used to study visual motion detection (Salazar-Gatzimas et al. 2016; Roy and van Steveninck 2016) to develop new correlated noise auditory stimuli that use increments and decrements in intensity to generate local correlations in intensity at specific offsets in frequency and time (see **Methods**). The correlations are local because they are limited to a single offset in frequency and time. We designed four stimuli with positive or negative correlations in intensity at an offset of 1/6 second, with the frequency offset directed either upward or downward by 1/15 octave (**Fig. 1b, S1**). These sounds were inharmonic, so that fundamental frequencies could not be used to judge pitch changes (McPherson and McDermott 2023, McPherson and McDermott 2018). We presented these stimuli to participants for 2 seconds and asked them to report whether they perceived the sound as having a rising or falling pitch profile over time.

Participants reported that upward-directed positive correlations (↑ +) rose in pitch over time, while those with downward-directed positive correlations (↓ +) fell in pitch over time (**Fig. 1c, Supp. Movie 1**). Being inharmonic, the stimulus did not carry F0 cues that could be leveraged to detect relative pitch. In these stimuli, the correlation over time reflected spectral features that partially persist for up to 1/3 of a second (see **Methods**). Thus, while this result suggests that humans can use spectrotemporal correlations to judge rising versus falling pitch, it is also conceivable that humans could track the transient spectral patterns in this stimulus over these short timescales.

Remarkably, however, when we presented stimuli with negative correlations in frequency and time, participants reported opposing percepts (**Fig. 1c, Supp. Movies 1 and 2**). That is, the upward-directed negative correlations (↑ –) sounded like they were falling in pitch, while the downward-directed negative correlations (↓ –) sounded like they were rising in pitch. Participants who consistently perceived rising or falling pitch in the stimuli with positive correlations also consistently perceived rising or falling pitch in the stimuli with negative correlations (**Fig. S1**). This striking illusion demonstrates that human audition is sensitive to negative spectrotemporal correlations. We emphasize that in these negative correlation stimuli, there do not exist spectral patterns that persist between frames, since negative correlations invert the transient, persistent spectral patterns. Thus, the inversion of percepts in this experiment is inconsistent with tracking-based models, and indicates that humans can use purely local spectrotemporal correlations to detect rising and falling pitch. Moreover, the inverted responses to negative spectrotemporal correlations are a direct analog to illusory reverse-phi visual motion percepts, which have been reported across many species and phyla (Clark et al. 2011; Orger et al. 2000; Anstis and Rogers 1975; Krekelberg and Albright 2005; Hassenstein and Reichardt 1956).

How does the strength of these spectrotemporal correlations relate to perception? To answer this question, we varied the coherence of the stimulus and again asked participants to judge whether tones were rising or falling in pitch (**Fig. 1d**). We titrated the coherence of the stimuli from 1 to 0 by randomly replacing correlated frequency-time elements with random ones, such that the coherence represented the fraction of original correlations remaining (see **Methods**). With high coherence, participants perceived rising and falling pitches in a pattern similar to the first experiment (**Fig. 1c**). As coherence decreased, however, the probability of judging a sound as rising tended towards chance (0.5). There were no significant differences between the curves for (↑ +) and (↓ –) or (↓ +) and (↑ –) (p > 0.05 for each, as measured by a two-way, repeated measures ANOVA). This indicates that inverting the stimulus correlation and direction led to indistinguishable percepts. These results reveal a clear monotonic relationship between the strength of spectrotemporal correlations and the strength of pitch change percepts, for both positive and negative correlations.

In vision, object tracking can integrate information between the two eyes, while correlation based algorithms rely on correlations within each eye (Lu and Sperling 1995). We next asked if spectrotemporal correlations for pitch motion detection are computed monaurally or binaurally. The structure of our correlated noise stimulus is created by summing a random binary mask with itself at a frequency-time offset (**Fig. 1e, Methods**). This allowed us to play one binary mask to the left ear and a shifted one to the right ear, so that neither ear alone was presented with any correlations. In this context, detecting spectrotemporal correlations can only proceed by integrating information across the two ears. We played all four types of binaural correlations to participants and asked them to judge whether they heard rising or falling tones. They reported the same pattern of percepts as in the monaural stimuli, though with average reported directions somewhat closer to chance. This demonstrates that the perception of rising or falling pitch can use information from both ears to integrate intensity information to compute spectrotemporal correlations. This is consistent with data showing that many cortical auditory neurons integrate signals from both ears (Kelly and Judge 1994), and represents a difference from correlation detection in the human visual system, which is predominantly monocular (Lu and Sperling 2001).

### Tuning of human spectrotemporal correlation detectors

Our next step was to characterize the spectral and temporal tuning of the correlation sensitivity we had observed. To do this, we designed a different kind of stimulus, one inspired by random dot kinetograms in visual neuroscience (Britten et al. 1992). In these stimuli, a medium intensity sound that played at all frequencies was interrupted by brief pips at different frequencies, 50 ms in duration (DeCharms et al. 1998). These pips either increased the intensity of a specific frequency or decreased it to zero (**Fig. 2a**, see **Methods**). After an initial set of pips were placed randomly in frequency and time, we added a second set of pips with a specific delay in time and change in frequency, yielding correlated “pip pairs.” These pairs had positive correlations when both pips were high intensity or both were silent, and negative correlations when one was high intensity and one was silent. This allowed us to create auditory stimuli with upward and downward-directed pairs of pips with positive or negative correlations (**Fig. 2b**). Like the stimuli used in **Fig. 1**, these stimuli had no auditory objects that persisted in time or frequency, but crucially, they allowed us to continuously vary the delay between correlated pips.

**Figure 2.**
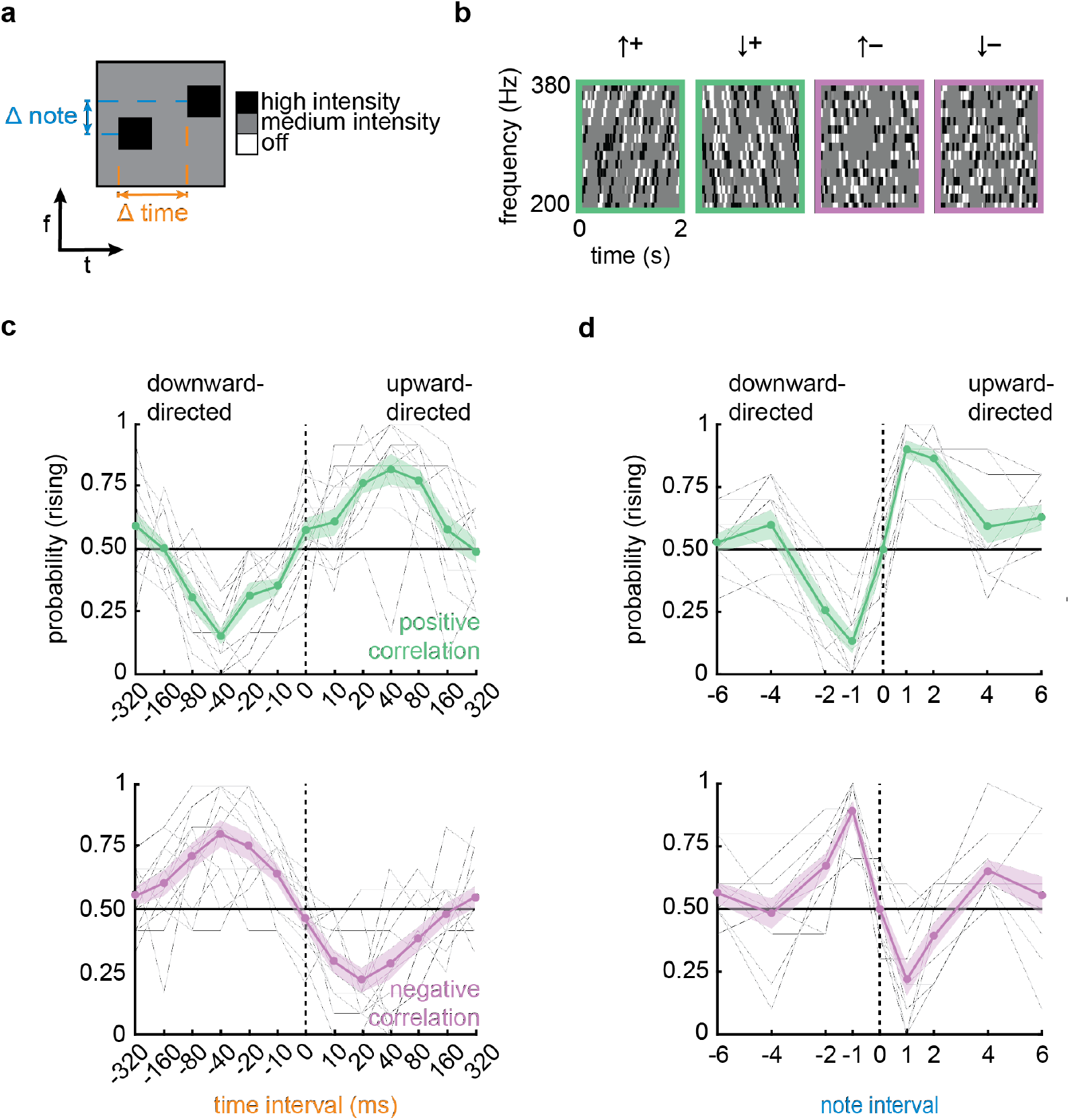
Correlation detection is tuned to small frequency changes and short delays in time. a) Diagram showing a correlated pip pair with a frequency displacement (Δ note) and a delay between pips. b) Spectrotemporal diagrams of 4 different correlated pip stimuli directed upward and downward with positive and negative correlations. Pip duration in these experiments was 50 ms. c) Perceived direction of stimuli with Δ note = +1 and varying pip delays; positive pip correlations (*top*) and negative pip correlations (*bottom*). One-way, repeated measures ANOVAs for the positive and negative correlation curves revealed significantly different responses across pip delays (all *p*s < 10^−21^). Gray lines are individual participant curves. Error shading represents ± SEM (N = 13). d) Perceived direction of stimuli using varying note intervals and 40 ms pip delays; positive pip correlations (*top*) and negative pip correlations (*bottom*). One-way, repeated measures ANOVAs for the positive and negative correlation curves revealed significantly different responses across note intervals (all *p*s < 10^−12^). Gray lines are individual participant curves. Error shading represents ± SEM (N = 13).

We first used these stimuli to map out the sensitivity to different delays between individuated tones. We kept the frequency change at 1/15 octave and swept values of the delay between correlated pips while asking participants to judge whether the pitch was rising or falling over time (**Fig. 2c**). For both negative and positive correlations and upward- and downward-directed displacements, we found that peak directional sensitivity occurred at a delay of around 40 ms. This peak did not change appreciably when the pip duration was shortened to 20 ms (**Fig. S2**). According to models for visual motion estimation, this peak sensitivity value reflects the typical relative delays in the circuits detecting local motion signals (Salazar-Gatzimas et al. 2016). The delay seen here is slightly longer than delays measured by similar experiments in human and fly visual systems (Bours et al. 2009; Salazar-Gatzimas et al. 2016). The timescale of sensitivity is typical of timescale differences over which asynchrony of tone presentation can be detected (Wojtczak et al. 2013).

We then measured sensitivity to the magnitude of displacements in frequency space. Using a similar method, we set the delay to 40 ms and varied the frequency displacement within each pip pair (**Fig. 2d**). We found that peak sensitivity occurred for tone displacements of 1/15th octave, though there was still significant direction-selectivity at 2/15th octave displacements (p < 0.05 for both positive and negative correlations by a paired t-test). This result shows that correlation-based motion detectors in the human auditory system are most sensitive to small shifts in frequency in the vicinity of 1/15th of an octave (4.7% changes in frequency) or less. This result is consistent with peak sensitivity for changes in complex sounds (Demany et al. 2009) and with the smaller values of frequency discrimination thresholds in humans (Sinnott and Aslin 1985).

### Sensitivity to spectrotemporal intensity patterns

Our positive and negative correlation stimuli each consist of multiple patterns in intensity over frequency and time. Upward-directed positive correlation (↑ +) stimuli consist of both high-high and low-low intensity combinations, whereas the negative versions (↑ –) consist of both high-low and low-high intensity combinations. Prior work using long-lasting spectrotemporal correlations in auditory stimuli has suggested that humans are selectively sensitive to the high-high combinations (Allik et al. 1989). Are humans sensitive to all four pairwise combinations, or to just a subset of them? To address this question, we generated new correlated pip auditory stimuli (**Fig. 3a**) where each stimulus had paired pips of only one of the four types: high-high, low-low, high-low, or low-high (see **Methods**). We asked participants to judge whether these different stimuli were rising or falling and recorded their responses (**Fig. 3b**). Participants were sensitive to all four different pairings with both upward and downward displacements.

**Figure 3.**
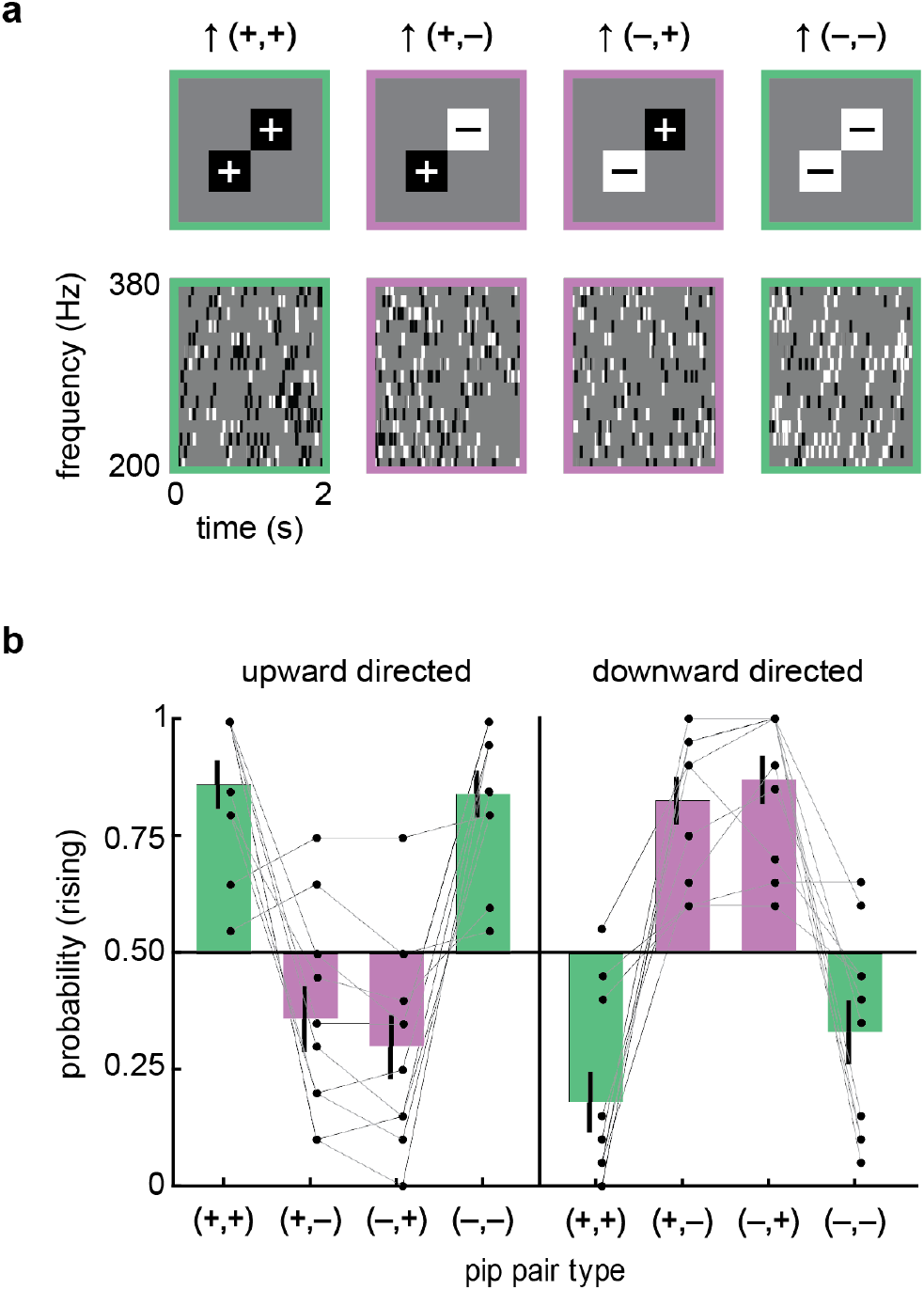
Sensitivity to all four pairwise intensity combinations contribute to rising and falling pitch perception. a) Frequency-time diagram of 4 different pip combinations, presented with 40 ms delays. b) Probability of perceiving rising pitch for each of the four intensity combinations directed upward (*left*) and downward (*right*). Paired t-tests comparing upward-versus downward-directed stimuli for each matched pair revealed significant direction selectivity across all pitch direction judgements (all *p*s < 10^−4^). Error bars represent mean ± SEM (N = 10).

In visual motion detection, one generalization beyond pairwise correlations involves so-called triplet correlations (Hu and Victor 2010; Fitzgerald et al. 2011). In vision, triplet correlation stimuli are patterns that contain spatiotemporal correlations over three points in space and time, but no pairwise correlations, and can elicit visual motion percepts in humans (Clark et al. 2014; Hu and Victor 2010), flies (Clark et al. 2014; Chen et al. 2018), and fish (Yildizoglu et al. 2020). Visual motion detection algorithms are sensitive to this higher-order correlative structure, but is the same true in audition? When participants were presented with auditory analogs of visual triplet correlation stimuli (see **Methods**), they did indeed perceive auditory motion (**Figure S3**) and did so in a pattern very similar to the pattern found in fly and fish visual perception. This correspondence across both species and modalities points to significant similarities in the neural algorithms used by animals in processing auditory (spectral) and visual (spatial) motion.

### Psychophysical and cortical signatures of opponent subtraction of spectral motion signals

When we presented positively and negatively correlated stimuli, we discovered a striking symmetry: Tuning of percepts of negative correlation stimuli matched the tuning of percepts of positive correlation stimuli that were displaced in the opposite direction (**Fig. 2**). This clear symmetry seemed suggestive of an opponent architecture. To investigate this, we first built a toy model of a motion energy model unit to describe a hypothetical directionally tuned auditory unit (**Fig. 4a**). The model unit temporally filtered and summed sound intensity at two adjacent frequencies in a pattern that enhanced upward-directed spectral motion, similar to prior suggestions (Brown and Cooke 1994), before sending the filtered signal through a quadratic nonlinearity (Adelson and Bergen 1985). This model was not intended to be a realistic model of neural processing, but rather a tractable simplification of spectrotemporal correlation detection algorithms (Adelson and Bergen 1985).

**Figure 4.**
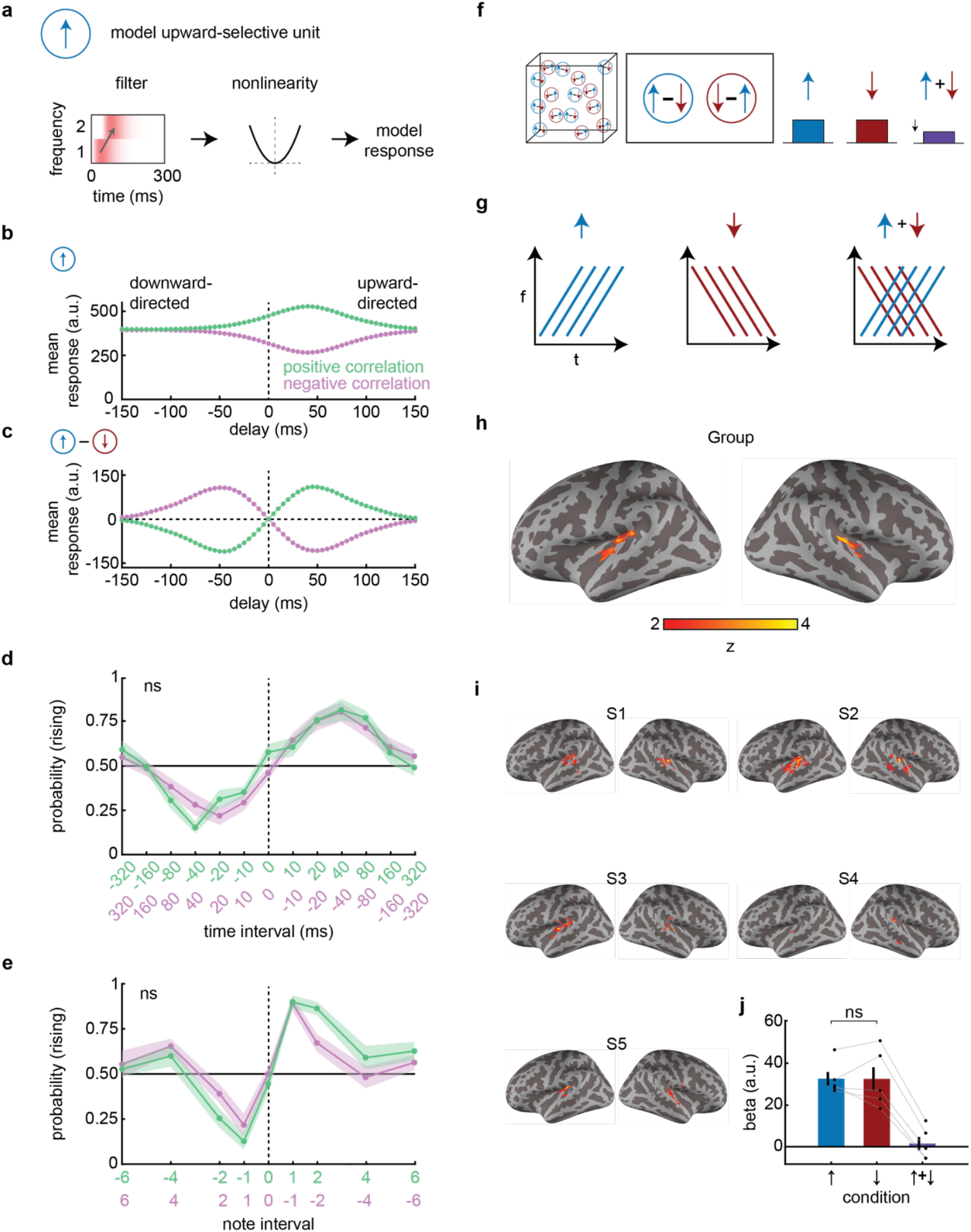
Bilateral regions of human auditory cortex show signatures of opponency. a) A simple model auditory unit that responds more to upward direction spectral motion than downward-directed spectral motion. The stimulus spectrogram is convolved with an upward-oriented spectrotemporal filter before the result is squared, as in a motion energy model (Adelson and Bergen 1985). b) Mean response of the unit to correlated pip stimuli with different delays and correlation signs, corresponding to upward and downward-directed positive and negative correlations. c) As in (b), but for an opponent signal, consisting of an upwardly tuned unit response minus an identical unit tuned to downward motion. d) Comparison of P(rising) for positive and negative correlation stimuli sweeping time interval, aligning upward-directed positive correlation stimuli with downward-directed negative correlation stimuli. Data replotted from Figure 2. The curves were not significantly different (p > 0.05 by a two-way, repeated measures ANOVA). e) As in (d) but for sweeping the tone difference. The curves were not significantly different (p > 0.05 by a two-way, repeated measures ANOVA). f) Conceptual schematic of opponency in brain regions. An opponent voxel/region would respond strongly to rising and falling tones but be suppressed by the sum of the two stimuli. g) Stimulus design. Stimuli were rising, falling, or summed rising and falling. h) Group level analysis. A bilateral region within auditory cortex responded less to summed stimuli than non-summed stimuli. Cluster-corrected with false positive rate at p < 0.05 with a cluster-forming threshold of 20 voxels. i) Individual level analysis. Regions in auditory cortex across subjects responded less to summed stimuli than non-summed stimuli. Cluster-corrected with false positive rate at p < 0.05 with a cluster-forming threshold of 20 voxels. j) Control analysis showing symmetric beta values in response to rising and falling stimuli in individually defined opponent ROIs (p > 0.05 via one-sample t-test). (Note that all beta values are relative to an implicit baseline that includes responses to ambient scanner noise.) Error bars represent mean ± SEM (N = 5).

When we presented this model with correlated pip stimuli (**Fig. 2**), it responded at an elevated baseline level, with deviations that depended on the direction and sign of the stimulus correlation (**Fig. 4b**). As designed, it responded more to upward-directed positive correlations than to downward-directed ones. Since this model relies solely on pairwise correlations, it was also expected that negative correlation stimuli elicited equal and opposite deviations to positive correlation stimuli (**Fig. 4b**). Crucially, however, in this model, negatively correlated stimuli exhibit a different tuning from oppositely directed positive stimuli; that is, inverting the correlation is not equivalent to inverting the correlated pip direction (i.e., the temporal delay in the pips). Thus, this model does not predict the symmetry we observed in the psychophysics (**Fig. 2**).

We next created an opponent signal by subtracting signals from two model units with opposite directional tuning (**Fig. 4c**). This opponent signal responded to positively correlated stimuli with positive and negative values when they were directed upward and downward (**Fig. 4c**, *green*). Critically, this opponent signal possesses an important symmetry: Responses to negatively correlated stimuli have the identical tuning as positively correlated stimuli in the opposite direction. Thus, upward-directed negative correlation stimuli yield identical responses as downward-directed positive correlation stimuli. We also derived this result analytically (see **Methods**): When motion energy signals are opponently subtracted, negative correlation stimuli elicit mean responses that match oppositely directed positive correlation stimuli.

To test whether our data contained this symmetry, we compared percepts of negative correlation stimuli to percepts of positive correlation stimuli in the opposite direction, for both frequency change and delay time tuning (**Fig. 4d, e**, replotting data from **Fig. 2**). The curves appeared to fully superimpose. ANOVA tests confirmed that there was no measurable difference between the positive correlation curves and the flipped negative correlation curves (see figure legends for statistics). This robust symmetry between positive and negative correlation stimuli has also been found in visual motion detection in fruit flies (Salazar-Gatzimas et al. 2016) and in humans (Bours et al. 2009).

In primate vision, opponent subtraction occurs in visual area V5, also called MT (Snowden et al. 1991; Qian and Andersen 1994), which is causally involved in visual motion percepts (Salzman et al. 1992). Similarly, fly visual systems also subtract motion signals with opposing preferred directions (Mauss et al. 2015). Motivated by our psychophysical results, by analogies with vision, by proposals for opponent subtraction in spectral direction (Demany and Ramos 2005), and by spectral direction opponent auditory cells found in bats (Andoni and Pollak 2011), we reasoned that human auditory cortex might possess signatures of opponent processing.

We followed the logic of previous functional magnetic resonance imaging (fMRI) studies that identified opponent signals in human cortical area MT by using visual stimuli that summed motion in opposite directions (Heeger et al. 1999). To start, we assume that cortical voxels involved in detecting spectral motion contain units that respond preferentially to rising tones and units that respond preferentially to falling tones, but not units that respond to both (**Fig. 4f**) (Tian and Rauschecker 2004). Such a voxel should thus respond reliably to stimuli containing either rising or falling tones. The key distinction here between a system with or without opponency lies in its response to a summed stimulus that contains superimposed rising and falling tones: If units are opponent, then the summed stimulus should cause a decrease in voxel activity due to a net suppression of signals in units with opponent responses (Heeger et al. 1999). We therefore designed simple stimuli consisting of rising tones, falling tones, or their sum (**Fig. 4g, S4**) and presented them to subjects while measuring blood-oxygen-level-dependent (BOLD) signals via fMRI.

We searched within a broad auditory cortex mask for voxels that responded more to the non-summed (rising or falling) stimuli than to the summed (opponent) stimulus (see **Methods**). Strikingly, at both group and individual levels, a bilateral region within superior temporal cortex was significantly more activated by the non-summed stimuli than by the summed stimulus (**Fig. 4h, i**), consistent with opponency. The group map extended over multiple bilateral functional subregions of the human auditory cortex (Glasser et al. 2016), including core regions A1 and RI, Area 52, and lateral and medial belt regions (**Fig. S4**). According to the opponency hypothesis, activity in opponent voxels should be similar in magnitude for rising and falling stimuli and suppressed for the summed stimulus. Thus, we wanted to ensure that our result followed this symmetry and was not biased by either the rising or falling stimulus alone (see **Methods**). Activity in putative opponent regions was indeed comparable for rising and falling tones (**Fig. 4j**). Overall, our fMRI findings demonstrate that a key result from our behavioral studies — the clear symmetry between positive and negative correlation percepts — lead to a specific neural hypothesis that was borne out in neuroimaging data. To our knowledge, these regions of human auditory cortex have not previously been identified as potential loci for opponent spectral motion signals, though they are broadly consistent with regions of auditory cortex sensitive to spectral motion (Schönwiesner and Zatorre 2009; Langers et al. 2003).

### Positive and negative correlation spectrotemporal cues signal tone modulation in speech

Is there an ecological advantage in detecting both positive and negative spectrotemporal correlations in intensity? To address this question, we asked how spectrotemporal correlations could help predict rising and falling frequencies in naturalistic sounds. This approach follows literature in visual motion detection, in which different spatiotemporal cues have been analyzed to understand how they can be used to infer motion in scenes (Fitzgerald et al. 2011; Potters and Bialek 1994; Fitzgerald and Clark 2015; Nitzany and Victor 2014). To examine rising and falling pitch in naturalistic sound, we chose to examine human speech, where tone modulation contains critical semantic information in both tonal and non-tonal languages (Hirst and Di Cristo 1998; Gandour 1983; Yip 2002). Since humans are sensitive to both positive and negative pairwise correlations in frequency and time, we hypothesized that these correlations could convey information about the direction and speed of tone modulation in human speech, in addition to the F0 cues also associated with human speech (Fujisaki 1983). Following in the tradition of relating auditory processing to natural sounds (Singh and Theunissen 2003), we analyzed corpora of spoken English and Mandarin and examined how rising and falling tone modulation is related to underlying positive and negative pairwise spectrotemporal correlations in intensity (**Fig. 5, Methods**).

**Figure 5.**
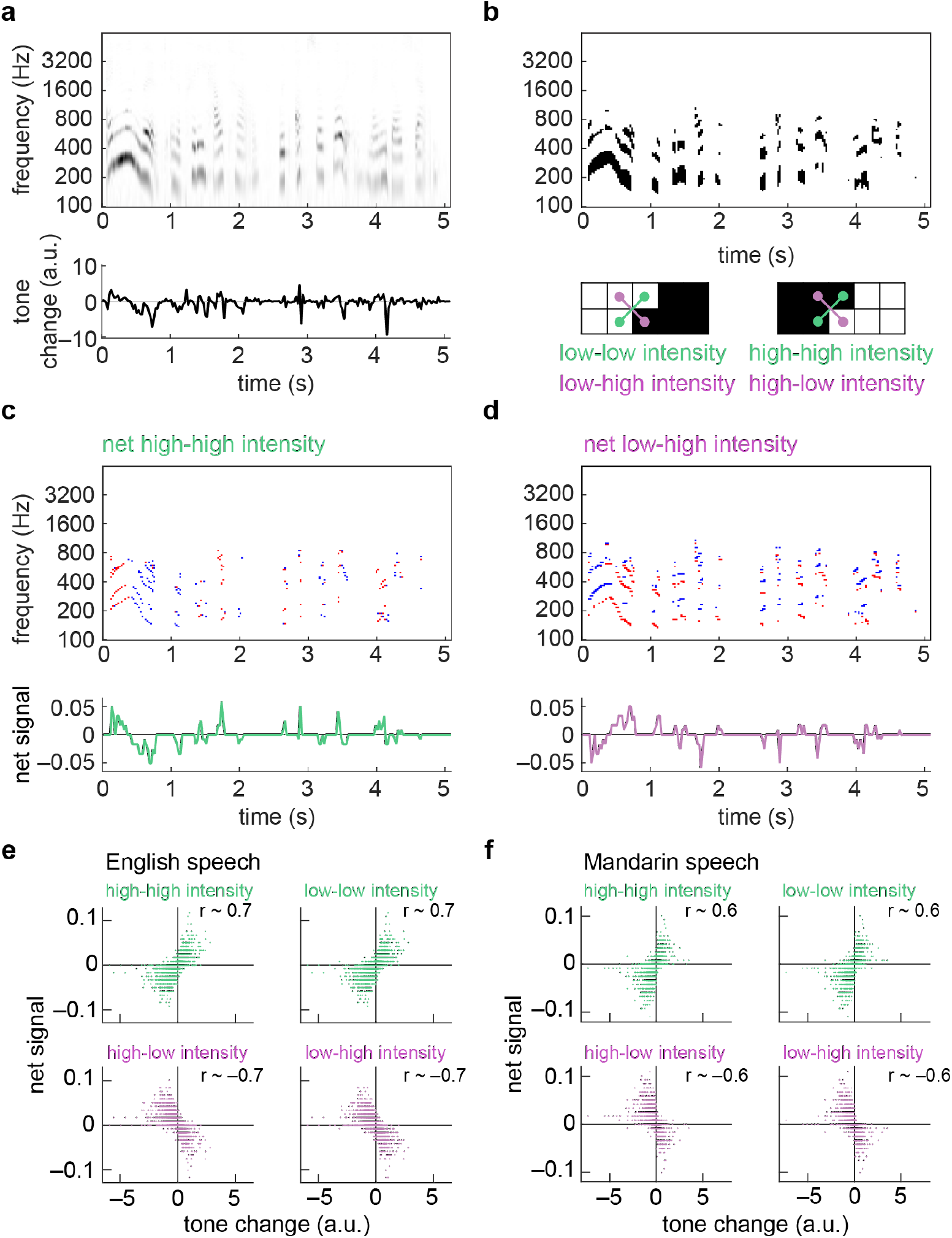
Rising and falling tone in spoken language can be detected through both positive and negative pairwise correlations. a) Spectrogram of voice saying, “Anyone lived in a pretty how town (with up so falling many bells down)” (*top*). Intonation velocity estimate from spectrogram (*bottom*, see **Methods**). Positive tone changes correspond to rising frequencies in the sound. b) Binarized spectrogram from (a) (*top*). Four distinct high and low intensity frequency-time combinations in the binarized spectrogram (*bottom*). c) Net high-high intensity instances at each frequency and time in the binarized spectrogram in (b) (*top*). Red is +1, blue is –1, white is 0. Frequency-averaged net high-high intensity signal (*bottom*). d) Net low-high intensity patterns at each frequency and time in the binarized spectrogram in (b) (*top*). Red is +1, blue is –1, white is 0. Frequency-averaged net low-high intensity signal (*bottom*). e) Correlations between the tone change estimate at each time and the frequency-averaged net signals for high-high, low-low, high-low, and low-high intensity patterns. Data from English speech corpus (Panayotov et al. 2015). f) As in (e) but for Mandarin speech corpus (MagicData 2019).

Our analysis took several steps, which were intended simply to analyze the structure of spectrotemporal correlations in speech, not reproduce any processing in the ear or downstream auditory system. First, we computed spectrograms for each of the speech recordings (**Fig. 5a**, *top*). We then used an optical flow algorithm to estimate how the tones in the sound changed at each point in time, a quantity we termed the tone change. The tone change represents the degree to which the sound was rising or falling in frequency at each time (**Fig. 5a**, *bottom*, see **Methods**). Next, we binarized the spectrogram and looked for specific patterns of intensity in frequency and time, examining all four patterns of high and low intensity combinations: high-high, low-low, high-low, and low-high intensity (**Fig. 5b**), which constitute spectrotemporal patterns that contain positive correlations (high-high and low-low intensity) and negative correlations (high-low and low-high intensity). We next computed the local net signal for each pattern at each frequency and time by subtracting the downward-directed patterns from the upward-directed ones for each of the four patterns (**Figs. 5c, d**). Finally, we averaged these local net signals over all frequencies to obtain a net pattern signal at each time point (**Figs. 5c, d**).

Computing the net pattern signals is an operation consistent with the opponency we observed psychophysically and in fMRI (**Fig. 4**). For the high-high intensity patterns, there was a positive correlation between the time trace of the net pattern signal and the tone change. For the high-low intensity patterns, the correlation was negative. This correspondence suggests that both positive and negative spectrotemporal correlations contain information about tone changes that could be useful to listeners in detecting rising and falling tones in speech.

To see whether this result generalized, we analyzed hundreds of speech snippets that totaled over 90 minutes in English and 40 minutes in Mandarin Chinese (**Fig. 5e, f**, see **Methods**). In English, the tone changes should be dominated by intonation, while in Mandarin Chinese, the tone changes should reflect both intonational and within-syllable changes in tone (Hirst and Di Cristo 1998; Gandour 1983; Yip 2002). We reproduced the analysis of the different intensity patterns, and then correlated the net signal for each pattern with the computed change in tone. In both English and Mandarin Chinese, the two positive correlation patterns (high-high and low-low intensity) produced a strong positive correlation (*r* > 0.5) with the tone change, whereas the two negative correlation patterns (high-low and low-high intensity) produced a strong negative correlation (*r* < – 0.5) (**Fig. 5e, f**). These results show that all four patterns could be useful in estimating tone changes in speech. The negative stimulus correlation produced an *anti*-correlation with tone changes, which explains why they elicit percepts in the opposite direction: upward-directed negative spectrotemporal correlations indicate downward-directed tone changes. We obtained similar results when we processed the speech recordings with continuous rather than digital operations to obtain positive and negative spectrotemporal correlations (see **Methods, Fig. S5**). Thus, this analysis provides an ecological explanation of the observed inverted percepts to negative auditory correlations: negative spectrotemporal correlations provide useful information for distinguishing between sounds with rising versus falling frequencies. The reversed perceptual direction for the negative correlation stimuli matches the relationship of negative correlations with rising and falling frequencies in sounds.

## Discussion

In the studies reported here, we have demonstrated that humans are sensitive to positive and negative spectrotemporal correlations in intensity over frequency and time as they discern whether a sound is rising or falling in pitch (**Figs. 1-3**). These correlations can be sensed over durations of less than 100 ms (**Fig. 2**). The perception of negative spectrotemporal correlations ruled out spectral pattern tracking-based explanations of our results; it also mirrors a powerful visual phenomenon, the reverse-phi illusion, in a different modality (audition) and over a different dimension of motion (frequency). Inspired by our behavioral results showing symmetry between inverting correlation and inverting direction, we hypothesized that the human auditory system might implement opponent subtraction, echoing a similar operation in visual motion detection. Using fMRI, we discovered that regions within human auditory cortex show signatures of opponency similar to those in visual cortical areas (**Fig. 4**). Finally, we demonstrated that both positive and negative spectrotemporal correlations can act as reliable cues to assess tone changes in speech (**Fig. 5**).

The stimuli we developed here (**Figs. 1-3**) in some ways resemble Shepard tones (Shepard 1964), which were designed to sound like they are unceasingly rising or falling. However, Shepard tones consist of periodic auditory features that persist over frequency and time (similar to **Fig. 4f**). Thus, the rising or falling of a Shepard tone could be assessed by simply tracking auditory features over time. The auditory stimuli we developed and investigated here, however, have no such persistent features –– a rising or falling percept must instead depend on detecting positive and negative pairwise spectrotemporal correlations within the stimulus. Thus, the strong percepts of rising and falling tones, which depended on the sign of the correlation, reflect an authentic auditory illusion in which there is no true rising or falling tone but only the imposition of specific local spectrotemporal correlations in intensity.

Sensitivity to spectrotemporal correlations in judging pitch direction likely acts in coordination with other algorithms for judging changes in relative pitch. In particular, changes in frequency can be judged over gaps of seconds (Demany et al. 2009), much longer than the correlations in our stimuli; this points to a different system for such judgements. Judgements about relative pitch tend to be made using fundamental frequencies in harmonic sounds, but also tracking of broader spectral patterns (McPherson and McDermott 2020, McPherson and McDermott 2023, McPherson and McDermott 2018). These examples suggest that auditory spectral motion processing is similar to visual motion processing in that there are a variety of likely distinct algorithms at play: changes can be detected by both local correlational algorithms and by slower, longer range “object-tracking” algorithms (Lu and Sperling 2001).

Our psychophysical findings inspired us to test the idea that opponent computations may be performed during spectrotemporal correlation detection. We found regions in both primary and non-primary auditory cortex across both Heschl’s gyrus and the superior temporal gyrus (STG) that may perform opponent computations to resolve net pitch direction (**Fig. 4h, i**). How might this relate to the neural underpinnings of speech perception? Human auditory cortex displays regional specialization, with areas that selectively encode different aspects of speech, primarily in the STG (Yi et al. 2019; Hickok and Poeppel 2007; de Heer et al. 2017; Li et al. 2021). Our results are broadly consistent with findings that regions within STG encode variability in speaker intonation and lexical tone (Tang et al. 2017; Li et al. 2021). Moreover, our observation that significant portions of Heschel’s gyrus also showed spectral motion sensitivity is broadly consistent with other work (Johnsrude et al. 2000), though we saw the effects bilaterally (**Fig. S4**) and, critically, with an opponent signature. Our results thus suggest that opponency may be a signature of pitch direction processing in circuits involved in simple pitch computations (in primary areas) and in more complex perceptual tasks like speech processing (in non-primary areas).

Canonical algorithmic models for motion detection are sensitive to negative correlations (Hassenstein and Reichardt 1956; Adelson and Bergen 1985), and more neurophysiologically-inspired models for motion detection are similarly sensitive to negative correlations (Zavatone-Veth et al. 2020; Mo and Koch 2003). At the single neuron level, units in rodent (Ye et al. 2010), bat (Andoni et al. 2007), and primate (DeCharms et al. 1998) auditory cortex display spectrotemporally oriented receptive fields, which should confer sensitivity to both positive and negative spectrotemporal correlations (**Fig. 4**) (Adelson and Bergen 1985). Our results suggest that neurons with this type of sensitivity could underlie spectrotemporal correlation detection in humans. Meanwhile, our psychophysical and fMRI results also suggest that units in multiple regions of auditory cortex exhibit directional opponency, a property observed in bat auditory neurons (Andoni and Pollak 2011). This direction opponency could arise in primary motion detectors (Badwan et al. 2019), or result from subtracting opposing cortical or subcortical motion signals (Kuo and Wu 2012; Lu et al. 2022).

There are well-established similarities in the processing of visual motion between invertebrates and vertebrates (Clark and Demb 2016; Sanes and Zipursky 2010; Borst and Helmstaedter 2015), phyla that diverged hundreds of millions of years ago. Our study shows that local correlational algorithms for motion detection also span modalities, since human audition and vision appear to employ similar computational motifs. Audition thus joins olfaction (Kadakia et al. 2022) as a non-visual sense where pairwise, local correlations can generate directional motion percepts. A critical aspect of our results is that sensitivity to pairwise stimulus correlations also includes sensitivity to negative correlations. This sensitivity to negative correlations is due in part to the mathematics of computing correlations (see **Methods**) (Adelson and Bergen 1985; Fitzgerald et al. 2011; Clark and Fitzgerald 2024), providing a conceptual framework for understanding the neural detection of motion that spans modality and species.

Lastly, negative correlations sensed in audition likely act as useful cues to infer real-world changes in the frequency domain (**Fig. 5**), just as they may aid in visual motion detection (Fitzgerald and Clark 2015; Salazar-Gatzimas et al. 2018). Thus, the illusory pitch motion described here is not just an interesting laboratory epiphenomenon. Rather, it reflects neural sensitivity to the statistics of the auditory world, with direct implications for everyday speech and music perception.

## Contributions

PAV and DAC designed auditory stimuli. PAV and SDM acquired data. PAV, SDM, and DAC analyzed and interpreted data. PAV, SDM, and DAC wrote the paper.

## Acknowledgements

This work was funded by a grant from the Wu Tsai Institute at Yale University. DAC and this work were funded by NIH R01 EY026555. We thank R. Aslin, E. V. Clark, H. H. Clark, I. Yildirim, and J. Zavatone-Veth for helpful discussions and comments on this project.

## Supplementary Figures and Movies

**Supp. Figure S1.**
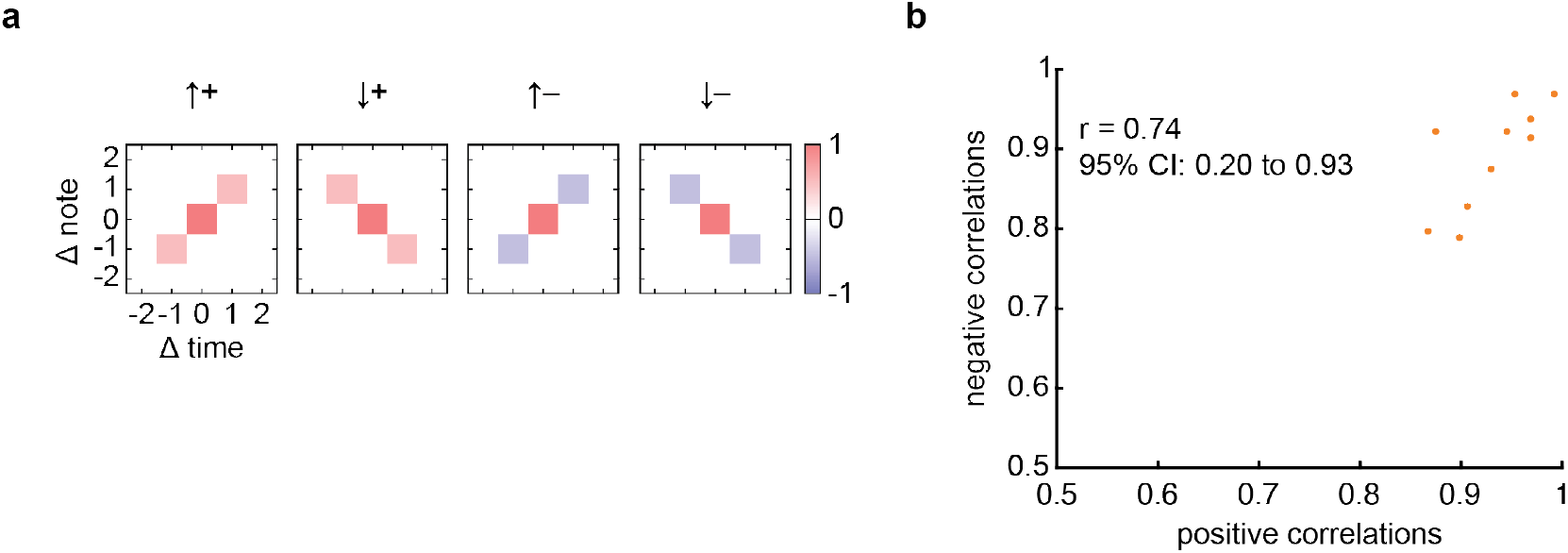
a) Stimulus autocorrelation plots at different note and time offsets for the stimuli in **Figure 1b**. The stimuli have positive or negative intensity correlations at a single spectrotemporal offset, directed either upward or downward in frequency over time. These plots are normalized so that the origin has correlation of 1. b) Correlation between perception of positively correlated and negatively correlated stimuli. To obtain the positive correlation values, we averaged P(rising) for the upward-directed, positive correlation stimuli with 1-P(rising) for the downward-directed, positive correlation stimuli. To obtain the negative correlation values, we averaged P(rising) for the downward-directed, negative correlation stimuli with 1-P(rising) for the upward-directed, negative correlation stimuli. Correlation coefficient is the Pearson correlation, and a 95% confidence interval is noted.

**Supp. Figure S2.**
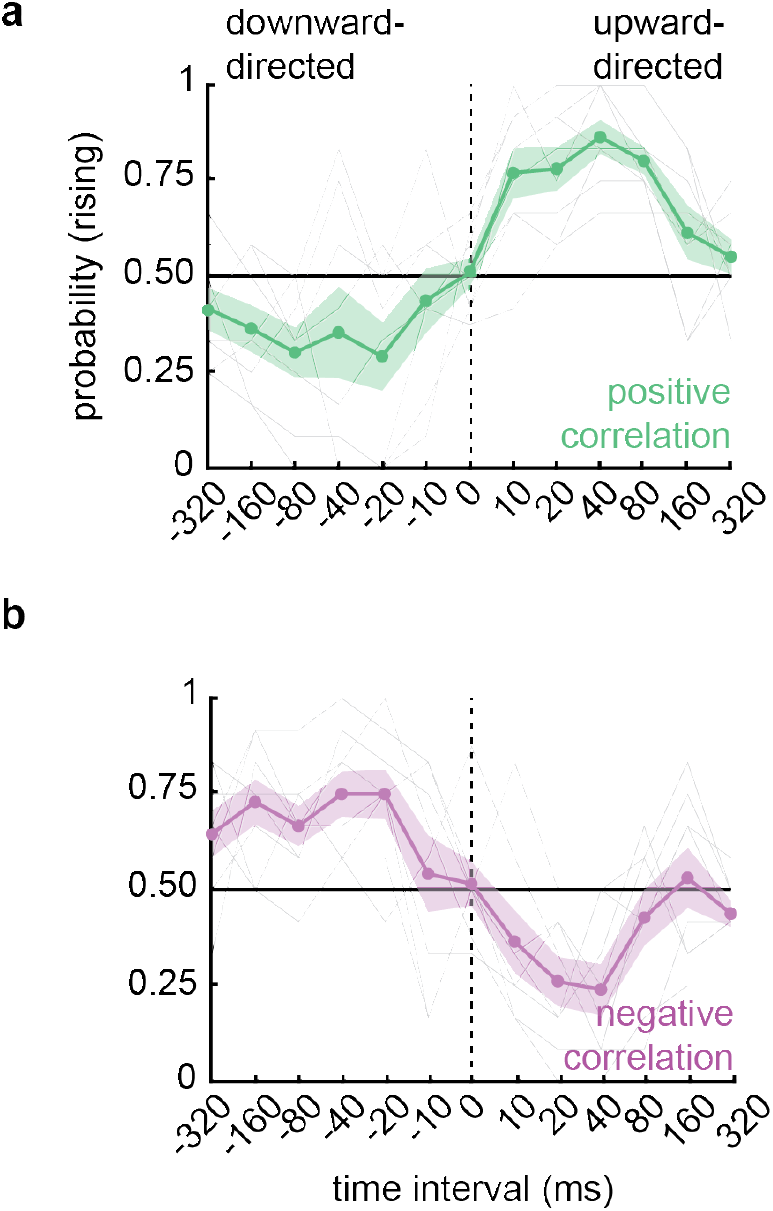
Interval sweep with a different pip duration. a) Perceived direction of positively correlated stimuli with varying pip delays and 20 ms pips. Sensitivity tends to peak around 40 ms delays, similar to the data in **Figure 2c**. A one-way, repeated measures ANOVA revealed significantly different responses across pip delays (p < 10^−10^). Error shading represents ± SEM (N = 9). b) Perceived direction of negatively correlated stimuli with varying pip delays and 20 ms pips. Sensitivity tends to peak around 40 ms delays, similar to the data in **Figure 2c**. A one-way, repeated measures ANOVA revealed significantly different responses across pip delays (p < 10^−8^). Error shading represents ± SEM (N = 9).

**Supp. Figure S3.**
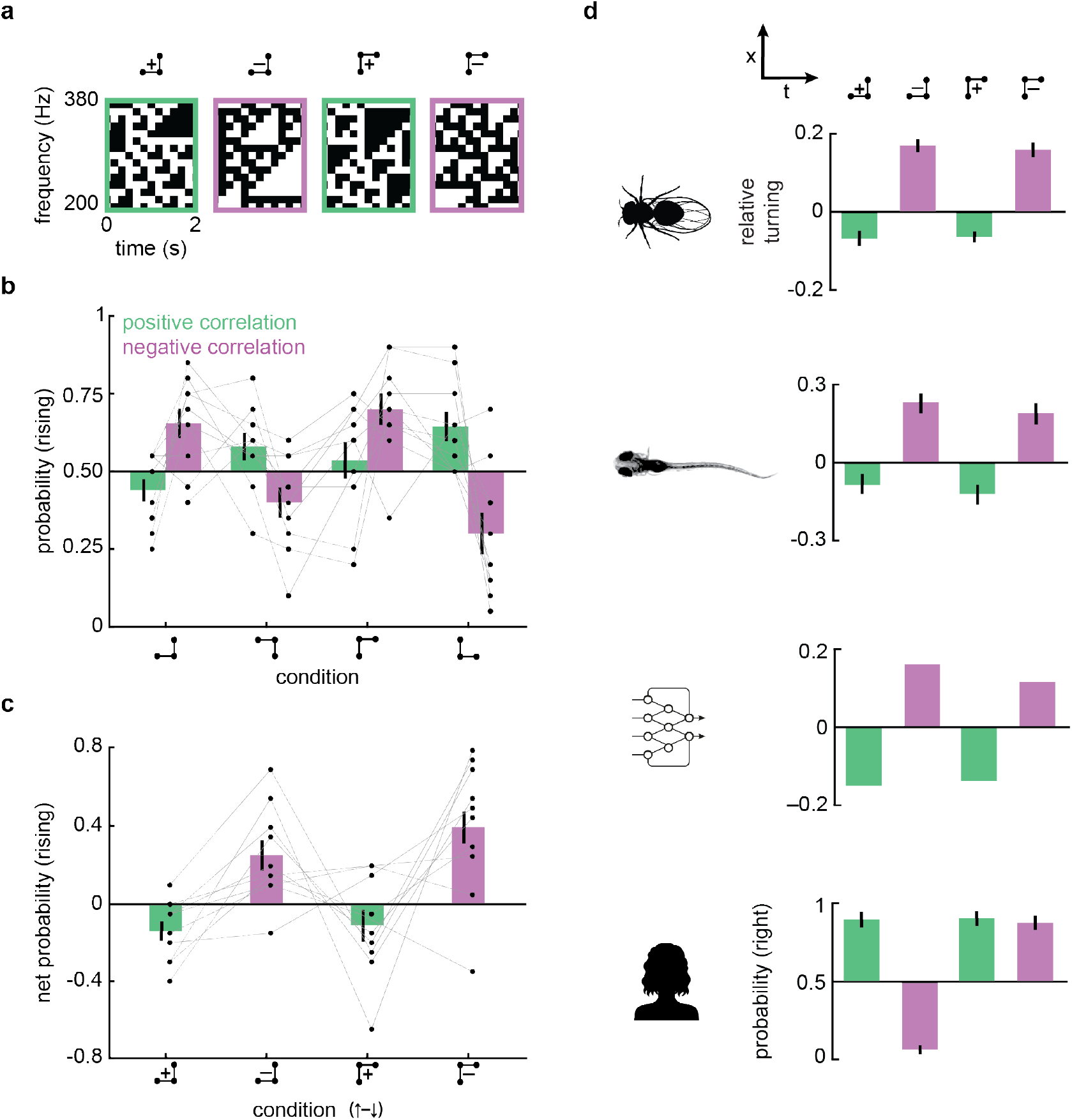
Human auditory sensitivity to 3-point glider stimuli resembles visual sensitivity in different species. a) Diagram of 3-point glider stimuli in frequency and time (Hu and Victor 2010). 3-point glider stimuli contain correlations between triplets of points as denoted by the barbell diagrams, and contain no pairwise correlations. Thus, motion percepts with these stimuli would have to rely on correlations beyond pairwise ones. b) Perceived direction of 3-point glider stimuli. Participants heard rising and falling tones in these triplet correlation stimuli. Error bars represent mean ± SEM (N = 10). c) Net perceived direction of 3-point glider stimuli with positive and negative correlations. The net probability rising is computed by subtracting the downward-directed P(rising) from the upward-directed P(rising) in panel (b). Positively correlated stimuli were perceived as falling, while negatively correlated stimuli were perceived as rising. Paired t-tests revealed significantly different responses to positively and negatively correlated diverging gliders, and to positively and negatively correlated converging gliders (all *p*s < 10^−3^). Error bars represent mean ± SEM (N = 10). d) Net perceived direction of 3-point glider stimuli across various visual systems. Data is replotted from prior publications for fruit flies (Clark et al. 2014), larval zebrafish (Yildizoglu et al. 2020), a machine learning algorithm (Fitzgerald and Clark 2015), and human visual psychophysics (Clark et al. 2014). Human auditory percepts resemble fruit fly and zebrafish visual percepts and machine learning responses, but not the human visual percepts.

**Supp. Figure S4.**
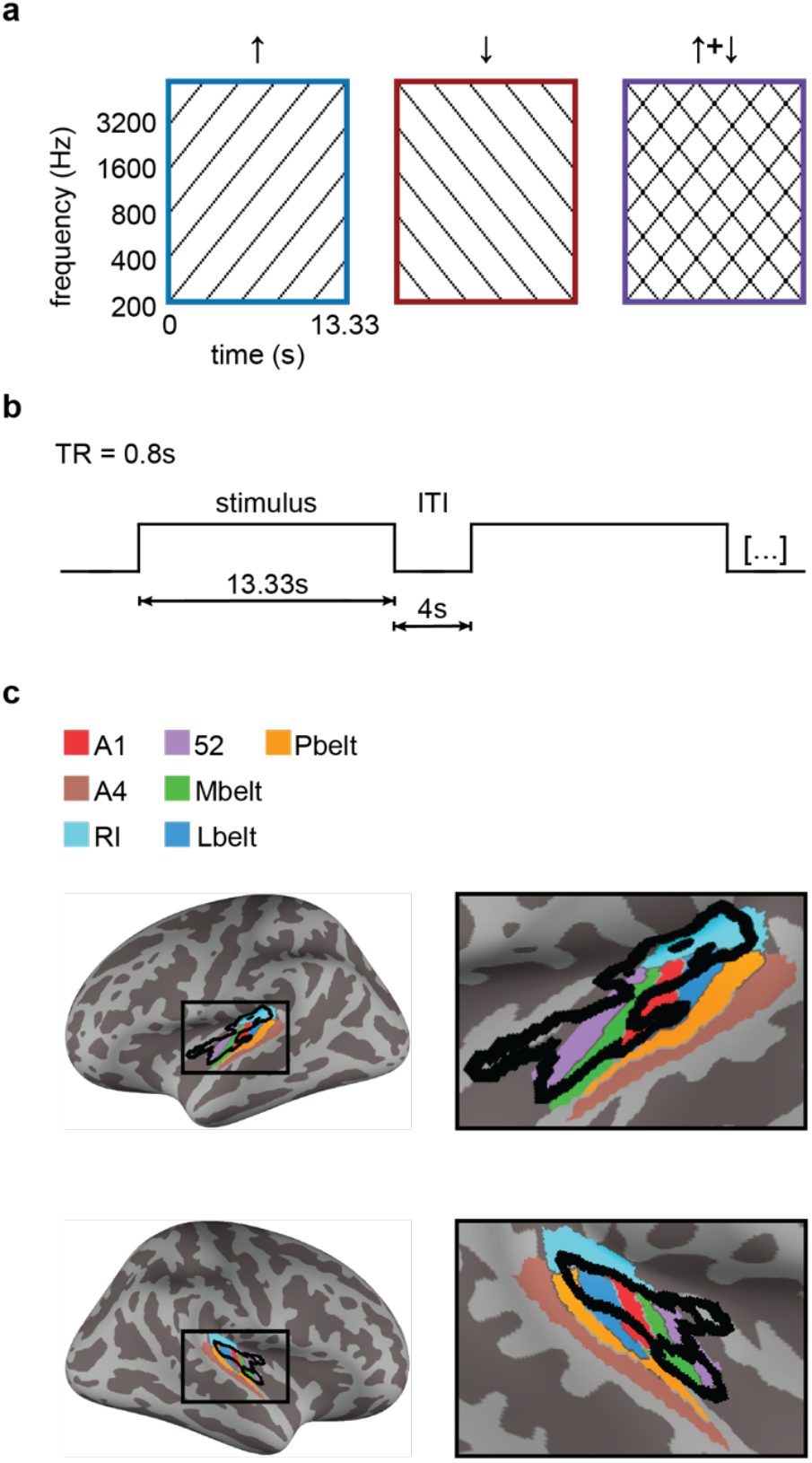
Further method and result details from fMRI opponency experiment. a) Depiction of actual stimuli used for the opponency experiment. b) Time course of fMRI trial structure. c) Group level analysis showing bilateral regions within auditory cortex that demonstrate significant opponent properties. Black outline reflects significant clusters from **Figure 4h**. Colored patches show cortical regions in accordance with (Glasser et al. 2016). *RI* = retroinsular cortex; *Mbelt* = medial belt of auditory cortex; *Lbelt* = lateral belt of auditory cortex; *Pbelt* = parabelt region.

**Supp. Figure S5.**
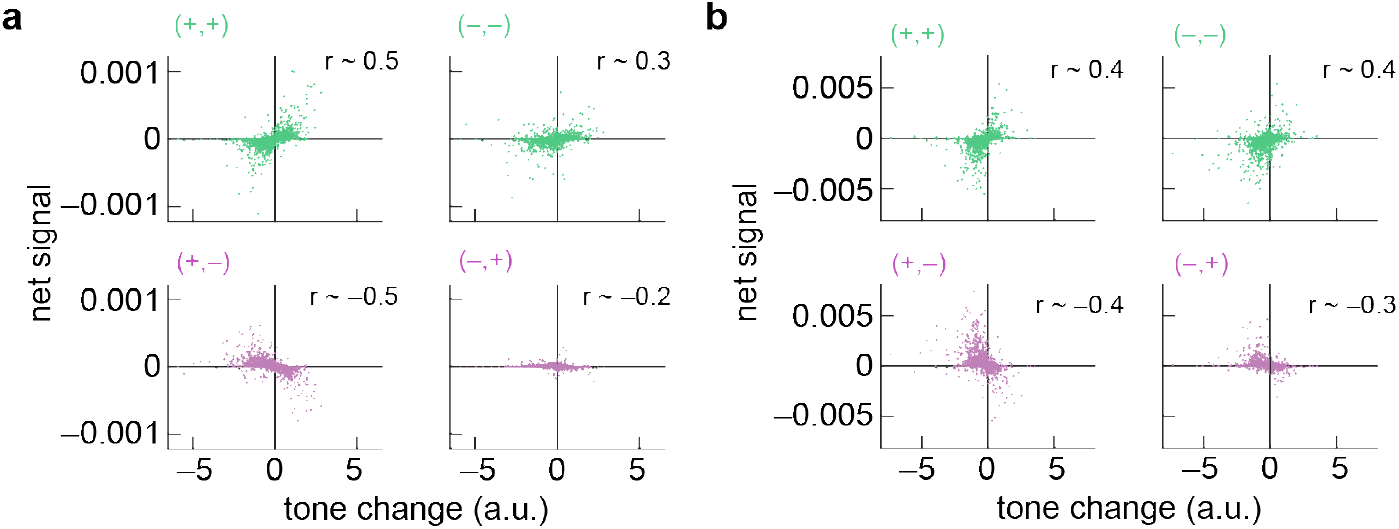
Multiplicative interactions of amplitude derivatives are informative about intonation direction. a) Correlations between the tone change estimate at each time and a continuous correlator model using only positive signals (+,+), only negative signals (–,–), and mixtures of the two (+,– and –,+) (see **Methods**). The correlations comprising the net signal were obtained by taking the derivative of the spectrogram amplitude in time, then multiplying derivatives of neighboring frequencies with a time-step delay and subtracting a mirror image product. Signals were rectified before multiplication to obtain the four pairs of multiplied signals, which together add up to a full correlator model. The net signals computed from (+,+) and (–,–) pairs correlated positively with tone change, while the net signals from (+,–) and (–,+) pairs correlated negatively with tone change. Data from English speech corpus (Panayotov et al. 2015). b) As in (a) but with data from Mandarin speech corpus (MagicData 2019).

**Supp. Movie 1**. Demonstration of positive and negative pairwise correlations using ternary correlated noise stimuli, as in Figure 1.

**Supp. Movie 2**. Demonstration of positive and negative pairwise correlations using ternary correlated noise stimuli, analogous to the stimuli in **Supp. Movie 1** but in visual motion detection (Salazar-Gatzimas et al. 2016).

## Methods

### Psychophysical measurements

All participants (N = 33; 12 female; mean age: 23.3 years, range of 18 years to 32 years) provided informed, written consent in accordance with procedures approved by the Yale University Institutional Review Board. To measure human psychophysical curves (**Figures 1-3**), we recruited participants with self-reported normal hearing from within the university population. Participants were seated in a quiet room, wearing headphones (Model DT 770 PRO, Beyerdynamic, Heilbronn, Germany) to listen to various sound stimuli and make perceptual judgments. The sounds were created in Matlab and presented using Psychtoolbox (Kleiner et al. 2007; Brainard 1997; Pelli 1997) on a Macbook Pro, using its native soundcard. Participants adjusted the intensity to a comfortable level. Each sound was played for 2 seconds, after which participants were cued to judge, to the best of their ability, whether it sounded like a rising or falling tone. To ensure they understood the task, participants went through several example sounds with the researcher before beginning the experiment. Participants usually completed two experiments lasting approximately 15 minutes each. The data was analyzed using custom code written in Matlab. The code to produce the sounds, all anonymized data, and the code used to analyze the data and produce Figure 1-3 are all publicly available at: [GitHub repository here, to be made available on publication].

### Creating correlated sounds

We created complex sounds containing multiple frequencies, following the design of visual stimuli that have been informative in that field. To do this, we created a comb of constant carrier frequencies, with frequencies ranging over 6 octaves from 200 Hz to 6400 Hz, with 15 frequencies per octave, equally spaced in log-space. The sampling frequency was chosen to be 20 kHz for all experiments. Each carrier frequency was then multiplied by a slower, time varying envelope, before the frequencies were summed to make the overall waveform for that sound.

Mathematically, the sound waveform, *w*(*t*), looks like:

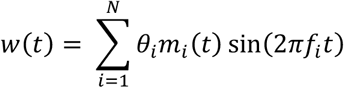

Where the *f*_*i*_ is the indexed carrier frequencies, *t* is sampled at 20 kHz, and the value *θ*_*i*_ Was chosen to roughly equalize the perceptual salience of the different frequencies, using the ISO standard 226 at 60 dB. (We note that in various tests in lab, this perceptual salience scaling was not critical for the percepts we measured; since we included it in initial experiments, we included it for all stimuli in this study.) It remains to compute the suite of *m*_*i*_(*t*) envelope functions to create each sound. The envelope functions were computed as outlined below. All envelope functions are computed to have non-negative binary or ternary values, and were filtered with a 25 ms low-pass filter in the ternary stimuli (**Fig. 1**) and at 0.5 ms low-pass filter in the pip stimuli (**Figs. 2, 3**) to eliminate sharp transitions. After all waveforms *w*(*t*) were created, they were scaled to have a minimum value of –1 and maximum value of +1.

#### Ternary pairwise correlations (**Figure 1b-d**)

To create sounds with only local, pairwise correlations between specific frequency and time offsets, we followed a protocol used in prior visual experiments (Roy and van Steveninck 2016; Salazar-Gatzimas et al. 2016; van Steveninck et al. 1996). Based on informal experiments attempting to optimize our own percepts, we discretized frequencies into 15 notes per octave and time into 1/6 second frames. This change in frequency is similar to the most salient change in frequency in a prior study (Demany et al. 2009). We then created an initial binary mask in this coarse-time representation, *B*_*ij*_, where *i* indexed the frequency and *j* the time step in 1/6 second intervals. In each trial, each element of *B* was chosen from a Bernoulli distribution with probability 0.5, then centered to have values of ±1/2 instead of 0 and 1. A ternary mask, *M*, was created by the following formula:

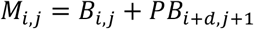

The mask is thus the binary matrix added back to itself with a displacement in frequency of *d* = ±1 for upward and downward-directed correlations. The mask is ternary, with values of 0, 1, and –1. The correlation parity is chosen by *P* = ±1, so that the offset matrices are added to create positive correlations and subtracted to create negative correlations. The discrete autocorrelation function of this mask *M* is equal to:

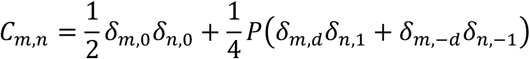

Where the δ_*i,j*_ terms are Kronicker delta functions (see **Fig. S1**). Importantly, the elements in the mask are not deterministically the same or different at the spectrotemporal offset of the correlated displacement, so that spectral patterns vary substantially at each temporal update of the stimulus.

A continuous time expression for the autocorrelation function is available in a prior work describing similar stimuli in vision (Roy and van Steveninck 2016).

The coarse-time matrix *M* was recentered to have values of 0, 0.5, and 1, then up-sampled to the sampling frequency *F*_*-*_ to create *m*_*i*_(*t*) at each frequency. The masks were filtered with a 25 ms low-pass filter to eliminate sharp transitions.

To create the stimuli with varying coherence, we replaced a fraction of mask elements with random ternary stimuli, drawn from the values (0, 0.5, 1) with probabilities (0.25, 0.5, 0.25). The fraction replaced was equal to (1 − *C*) where *C* is the coherence value.

#### Binaural pairwise correlations (**Figure 1d, e**)

To play sounds such that correlations only existed by integrating across the ears, we simply played *B*_*i,j*_ in one ear and *PB*_*i*+*d,j*+1_ in the other ear, for the correlations as described above to create the ternary pairwise correlations. To play these binary masks, we created two masks *M*_*i,j*_ = *B*_*i,j*_ and *M*_*i,j*_ = *PB*_*i*±*d,j*+1_ to play to the two ears. The matrices were recentered to have values of 0 and 1, then up-sampled to the sampling frequency. The masks were filtered with a 0.5 ms low-pass filter to eliminate sharp transitions.

#### Correlated pips with time and frequency offsets (**Figure 2**)

To create the correlated pip stimulus, we discretized frequency space into 15 tones per octave. We first initialized our masks *m*_*i*_(*t*) to be 0 for all times, sampled at the sampling frequency *F*_*-*_. We then placed initial delta-function pips in a Poisson distribution across all frequencies and times in our sound, at a rate of 4 pips per frequency per second. Positive and negative pips were equally probable, represented by mask values of ±1. We then created a second set of pips offset by the selected change in frequency and delay time, according to the two different correlation types. After imposing the correlations, the overall pip rate became 8 pips per frequency per second. We then convolved this event-trace with a boxcar function with the length of the pip duration to create the mask at *F*_*s*_. Pips had a duration of 40 ms in Figure 2 and 20 ms in Figure S2. Last, the masks were linearly transformed to be between 0 and 1 and filtered with a 0.5 ms low-pass filter to eliminate sharp transitions. The high intensity values corresponded to values of 1 in the mask, the low intensity to values of 0, and the background to values of 0.5.

#### Correlations between high and low intensity pips (**Figure 3**)

These stimuli were generated similarly to the correlated pips stimulus above. However, only two thirds of all pips were in correlated pairs of high-high, low-low, high-low, or low-high. In the case of the high-high correlated pips, the remaining third of pips consisted of randomly placed low intensity pips. In the case of low-low correlated pips, the remaining third of pips consisted of randomly placed high intensity pips. And in the cases of low-high and high-low, the remaining third were equally distributed between low and high intensity pips. Thus, the four types had equal numbers correlated pairs in each stimulus. The overall rate of pips for all stimuli was 6 pips per frequency per second.

#### Triplet correlations (**Figure S3**)

We made triplet correlation binary masks, discretized in frequency at time, following prior procedures (Hu and Victor 2010; Clark et al. 2014). The frequency was discretized in 15 tones per octave and time was discretized into 1/6 second frames. The frequencies began at 200 Hz and ranged over 5 octaves. The masks *m*_*i*_(*t*) were linearly transformed to have values of 0 and 1 and were filtered in time with a 0.5 ms low-pass filter to eliminate sharp transitions.

#### Rising, falling, and opponent tones (**Figure 5**)

To create the rising, falling, and opponent tones used in our fMRI experiment, we used frequencies discretized into 1/16 octave steps and time discretized into 1/6 second steps. Ascending tones were created from a binary mask equal to an ascending line of time-frequency elements in this discretized space (**Fig. S5**) and descending tones consisted of a descending line of time-frequency elements. The summed ascending plus descending was the sum of the two masks. All masks were filtered in time with a 0.5 ms low-pass filter to eliminate sharp transitions. We switched to 16 steps per octave for this experiment so that the ascending and descending stimuli never played the same frequency simultaneously, making the addition of the stimuli more straightforward.

Code to generate the sounds used in these experiments is available at [GitHub repository on publication].

### Model motion energy unit (Figure 4)

We created a toy model motion energy unit by convolving a linear filter with a sound spectrogram, then squaring the result. This model was simply intended to provide intuition about how spectrotemporal intensity signals could be processed — it is not intended to mimic the spectral processing of the cochlea and subsequent processing steps. That is:

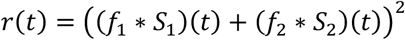

The filters were chosen to be:

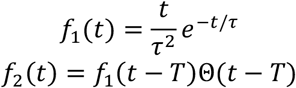

Where *f*_2_ is just a time-shifted version of *f*_1_ with a time shift of *T* = 40 ms. The function Θ is a Heaviside step function. The two filters are applied to adjacent frequencies in the spectrogram, *S*_1_(*t*) and *S*_2_(*t*), so that the filter enhances signals directed upward over time.

We computed the mean of *r*(*t*) over time to get the mean response for a given stimuli. Stimuli were created to match the correlated pip-style stimuli in Figure 2. The opponent response was computed as

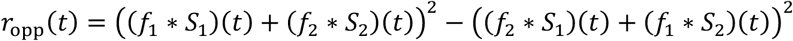

The second, negative term is the same as the first term but with the filter flipped in frequency space, so that it corresponds to a downward selective unit. This response was likewise averaged over time to produce the plots in Figure 4.

Matlab code to create **Figures 4b, c** is available at [Github repository on publication].

### Speech analysis

Spoken language databases were analyzed to ask how spectrotemporal correlations could act as indicators for rising and falling tones in speech. The analysis simply examines the spectrotemporal correlations in the sounds, rather than attempting to mimic any realistic auditory processing. Using Matlab, we first loaded short snippets of speech from two databases: 438 snippets constituting a total of 91 minutes of data from Librispeech, a corpus of read English (Panayotov et al. 2015); and 749 snippets constituting a total of 52 minutes of data from Magicdata Mandarin Chinese Read Speech Corpus (MagicData 2019), a corpus of read Mandarin. We computed a spectrogram for each snippet of speech using the Matlab command `spectrogram`; we extracted the spectral amplitude at a resolution of 40 samples per second with no overlap between samples, at 20 evenly spaced frequencies per octave from 100 Hz to 6400 Hz (**Figure 5a**). We estimated the rising/falling intonation change of the sound at each point using the Matlab command `opticalFlowHS`, which uses the Horn-Schunck method (Horn and Schunck 1981) to estimate directional local flow (typically optic flow) between frames. We averaged the calculated flow over frequencies to compute an estimate of the frequency “flow” with arbitrary units, which we termed tone change (**Figure 5a**). This method does not make strong assumptions about how changes in speech tone or frequency should be computed. It should work to extract tone changes from most complex sounds. We then examined estimators of this tone change as follows:

1. To compute binary correlations in frequency and time, we first binarized the spectrogram using Otsu’s method (Matlab command `imbinarize`) (Otsu 1975), which maximizes the variance between the binarized time-frequency element amplitudes while minimizing variance within each of the two categories (**Figure 5b**). We made 8 new binary frequency-time data arrays, containing Boolean values at each point in time and frequency, *V*_*t,f*,↑,±,±_ ≔ ({*A*_*t,f*_, *A*_*t*+1,*f*+1_} = {±1, ±1}) and an equivalent one for downward-directed intensity patterns. These matrices are records of the existence of each pattern of sound intensity at each time and frequency. From these, we computed the net signal of each pattern at each frequency by subtracting the downward-directed matrix from the upward-directed one. We last found the mean net signal over all frequencies for each pattern (**Figure 5c, d**). We computed the correlation between these mean net signals at each time point with the calculated upward or downward flow velocity (**Figure 5e, f**). Note that the sum of these net pattern signals sum to 0 over the four different patterns (±,±), so that the 4 signals are not independent.
2. To generate non-binarized correlation plots, we first linearly filtered the spectrogram amplitudes, *A*_*t,f*_, to take temporal derivatives: *F*_*t,f*_ = *A*_*t,f*_ − *A*_*t*,1,*f*_. We then used these derivatives, *F*_*t,f*_, which have positive and negative values, as inputs to a Hassenstein-Reichardt correlator model (Hassenstein and Reichardt 1956, Fitzgerald and Clark 2015). We then computed the net (+,+) correlations, for instance, as *N*_*t,f*,+,+_ = [*F*_*t,f*_] [*F*_*t*+1,*f*+1_]+− [*F*_*t*+1,*f*_] [*F*_*t,f*+1_]+, where [*x*]_+_ = *x* when *x* > 0 and [*x*]_+_ = 0 otherwise. A similar process computed the net (–,–), (+,–) and (–,+) correlations. We averaged these signals over frequency to obtain a single indicator of velocity at each point in time. These indicators were then correlated with the estimated tone change of the sound snippet at that point in time (**Figure S5**).

Code to analyze the spoken language databases and produce the panels in **Figure 5** is available at [GitHub repository].

### fMRI recordings and analysis

Whole-brain imaging was performed at the Brain Imaging Center at Yale University, on a Siemens 3 T Prisma MRI scanner using a 32-channel head coil. Functional data were acquired with a gradient-echo echoplanar pulse sequence (TR = 0.80 s, TE = 30 ms, flip angle = 52°, voxel size = 2.4 mm × 2.4 mm × 2.4 mm, MB acc. factor = 6). T1-weighted MP-RAGE anatomical images were collected as well (TR = 2.5 s, TE = 2.0 ms, flip angle = 8°, 208 slices, voxel size = 1.0 mm isotropic). Functional imaging in our sample (N=5; 1 female; mean age: 26.2 years; authors PAV and SDM were participants in the fMRI study) was performed in ∼5-minute runs, with the total number of functional runs per participant ranging from 3-5. Fifteen auditory stimuli were presented per run in an event-related design (5 each of three stimulus types: rising, falling, and summed). (We found that the correlated intensity sounds played in **Figure 1**, for instance, were not easily distinguishable by subjects in the scanner, so we did not examine neural responses to them in this study.) Each stimulus lasted for 13.33 s, separated by an inter-trial interval (ITI) of 4 s. The order of the three stimulus types was randomized in each run. Participants passively listened to the tones and were not required to render any responses. MRI-optimized noise-canceling headphones (Optoacoustics OptoACTIVE III) were used to limit effects of background scanner noise and the noise-cancelling software was trained on the EPI sequence sound features before each session using a brief calibration run.

The fMRI-Prep toolbox was used for preprocessing (Esteban et al. 2019). The anatomical image was corrected for intensity non-uniformity (INU) with N4BiasFieldCorrection (Tustison et al. 2010) and used as T1w-reference. The T1w-reference was then skull-stripped with a Nipype implementation of the antsBrainExtraction.sh workflow in ANTs, and tissue segmentation of cerebrospinal fluid (CSF), white-matter (WM), and gray-matter (GM) was performed on the brain-extracted T1w using FFAST (FSL 6.0.5) (Zhang et al. 2001). Volume-based spatial normalization to standard (MNI) space was performed through nonlinear registration with antsRegistration (ANTs 2.3.3). For each of the BOLD runs, a reference volume and its skull-stripped version were generated using a custom methodology of fMRIPrep. Head-motion parameters were estimated using MCFLIRT (FSL 6.0.5) (Jenkinson et al. 2002) and BOLD time-series were resampled into native space by applying the transforms to correct for head-motion, and the BOLD reference was co-registered to the anatomical reference using mri_coreg (FreeSurfer) followed by FLIRT. Co-registration was configured with 6 DOF. Several confounding time-series were calculated based on the preprocessed BOLD: framewise displacement (FD), DVARS and three region-wise global signals. The BOLD time-series were resampled into standard space, and volumetric resamplings were performed using ANTs.

Our main analyses involved constructing general linear models (GLMs) to quantify the effects of the three stimulus types within auditory cortex. GLM analyses were performed using Nilearn (Abraham et al. 2014). Confound regressors of no interest (generated using fMRIPrep, see above) were entered into each GLM. These included six standard motion regressors, the framewise displacement time course, and white matter and global signal time courses. Each stimulus type (rising, falling, and summed) was modeled using boxcar regressors over the entire stimulus presentation phase (13.33 s) of the relevant trials, and was convolved with the canonical double-gamma hemodynamic response function. The main contrast of interest at the group and individual levels compared BOLD responses to the non-summed directional stimuli (i.e., rising and falling) to the summed stimuli (i.e., superimposed rising + falling). The contrast was designed to highlight deviations from a null hypothesis of equivalent responses between directional and opponent stimuli. Individual subject runs were combined in a fixed effects analysis and then brought to the group level for mixed-effect analyses, where we controlled the false positive rate at p < 0.05 with a cluster-forming threshold of 20 voxels. Critically, individual-level results for all subjects were also analyzed and displayed, using the same thresholding parameters. All contrasts were performed within an *a priori* anatomical mask that consisted of any voxels crossing the 50% probability threshold within a combined bilateral probabilistic atlas (Harvard-Oxford) that included both the STG and Heschel’s gyrus. Individual and group results were projected onto the standard (MNI) cortical surface (FreeSurfer) for visualization.

A simple control analysis was also performed to ensure that the non-summed > summed results were not driven by a single non-summed stimulus (e.g., rising or falling) having a proportionally larger response, but rather by symmetric responses to the rising and falling stimuli. To perform this control analysis, we first extracted individualized regions of interest (ROIs) from the non-summed > summed contrast (using the threshold described above), and then extracted average beta values within that ROI for each stimulus type. We note that while this was of course not an unbiased ROI relative to the hypothesis that non-summed stimuli would on average show stronger activity than summed, it was unbiased relative to the hypothesis of symmetric responses to rising versus falling tones.

### Opponency implies a symmetry in responses with opposite correlations in opposite directions

The motion energy model uses pairwise correlations to extract motion information from input stimuli and seems to accurately represent important aspects of cellular physiology (Adelson and Bergen 1985). In the motion energy model, stimuli over space and time, *S*(*x, t*), are convolved with a space-time oriented linear filter, *H*(*x, t*). (In this section, we will derive results in space, but a frequency variable *f* could substitute for *x* and this approach would apply sound intensity over frequency rather than light intensity over space.) The result of the convolution is squared to obtain a response:

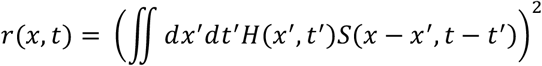

This response is stronger, on average, to stimuli with motion in the preferred direction than in the null direction. The preferred direction corresponds to the orientation of the filter *H* in space time, which amplifies signals when the motion direction aligns with the filter orientation. When the response is averaged over time and space, it yields a pleasing form in Fourier space, such that the mean response is the dot product of the stimulus power with a weighting function (Adelson and Bergen 1985):

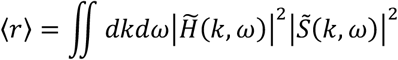

Where 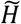 and 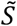 are the Fourier transforms of *H* and *S*. Therefore, to understand responses of this model, it is useful to compute the power spectrum of the stimulus.

For a random dot kinetogram in which the dots are displaced by Δ*x* in space and Δ*t* in time, the covariance density, *C*, of the stimulus is a function of the offsets in time and space, *x* and *t*:

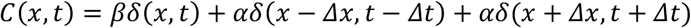

Where the first term is the stimulus autocovariance and the remaining two terms correspond to correlations in the stimulus at offsets of (Δ*x*, Δ*t*) and (−Δ*x*, −Δ*t*). For random dot kinetograms, β < 1 and α can take on positive or negative values for positively and negative correlated random dot kinetograms. This derivation is in continuous space, using Dirac delta function correlations; a similar result with discrete time and frequencies was found earlier in the methods for the ternary stimuli. The power spectrum of the stimulus is the Fourier transform of this covariance function:

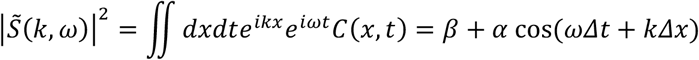

The power is highest/lowest along lines of constant phase in cosine, or when *ω*Δ*t* + *k*Δ*x* = *nπ*. When the *α* is negative, for negative correlation stimuli, this effectively changes the phase of the cosine by 180 degrees. The motion energy model says the mean response to such a stimulus, for a unit with filter *H*, is:

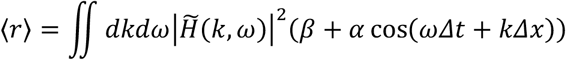

This is the type of curve shown in **Figure 4b**, in which there is a baseline response determined by *β* and the integral of 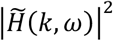. There is a modulatory term that depends on *α* and the dot product of 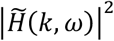 with cos(*ω*Δ*t* + *k*Δ*x*), which gives the modulation a directional tuning. This form means that the modulation inverts when the sign of the correlation (sign of *α*) inverts. If there is a peak response to a stimulus with correlation *α* at a specific Δ*t* and Δ*x*, then the peak will be equal and opposite when *α* is inverted. Importantly, however, the peak is not the same when the direction of the stimulus is inverted, that is when Δ*x* → −Δ*x*.

However, if we compute an opponent response, in which we subtract the response with one filter orientation from the response with the opposite filter orientation (inverting the *k* in the Fourier domain), then we find:

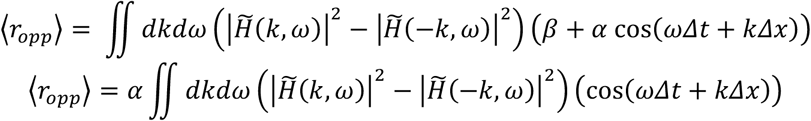

Here, we see that the opponent subtraction causes the *β* term to drop out entirely so that the remaining term is just proportional to *α*, the correlation in the stimulus. The mean opponent response can be computed for correlation stimuli with parameters *α*, Δ*t*, and Δ*x*: ⟨*r*_*opp*_(*α*, Δ*t*, Δ*x*)⟩. Because of the directional opponency, the response inverts when the stimulus is reversed in space:

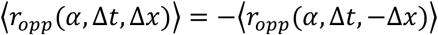

And because of the proportionality with the correlation, the response inverts when the stimulus correlation is inverted:

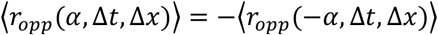

Therefore, for an opponent signal, inverting the correlation is equivalent to inverting the direction of the signal:

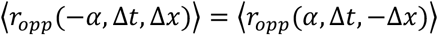

For any set of filters, as long as they are opponently subtracted, inverting the sign of the correlation is identical to inverting the direction of the stimulus, when computing the spatiotemporal average response. So when stimuli can be generated that have autocovariance structures like those in the ternary scintillator (**Fig. 1**) or in a random dot kinetogram (**Fig. 2**), if the computation is based on pairwise correlations and is opponent, the equations above show that the response will always be inverted when the stimulus correlation is inverted, and always be equivalent to inverting the direction of the stimulus. Therefore, opponency implies the sort of inversion symmetries we observed in our data, where inverting the correlation sign generates percepts with the same tuning as inverting the direction of the stimulus (**Fig. 4d, e**, but also visible in **Figs. 1-3**). Opponency also implies the sort of consistent symmetries between positive and negative correlation stimuli observed in human motion perception (Bours et al. 2009). We note that it is also possible to achieve this kind of symmetry using precisely defined filters that lead to opponent properties in single units, without a subtractive step (Badwan et al. 2019).

